# Cryo-EM Structure of R-loop monoclonal antibody S9.6 in recognizing RNA:DNA hybrids

**DOI:** 10.1101/2022.04.04.487086

**Authors:** Qin Li, Chao Lin, Haitao Li, Xueming Li, Qianwen Sun

**Affiliations:** Center for Plant Biology and, School of Life Sciences, Tsinghua University, Beijing 100084, China; Beijing Advanced Innovation Center for Structural Biology, Tsinghua University, Beijing, 100084, China; Tsinghua-Peking Center for Life Sciences, Beijing 100084, China

**Keywords:** R-loop, S9.6, RNA:DNA hybrid

## Abstract

The three-stranded chromatin structure R-loop is commonly found in the genomes of different species, and functions as the double-edged sword in gene expression and genome stability. The monoclonal antibody S9.6 specifically recognizes RNA:DNA hybrids, and has been widely used as a powerful tool to detect R-loops genome-widely. However, the structure basis and the molecular recognition mechanism of S9.6 to the nucleic acid substrates is still limited. Here, applying cryo-electron microscopy, we have determined a 4.9 Å structure of S9.6 antigen-binding fragment (Fab) complexed with RNA:DNA hybrids. We found that the native Fab cleaved from S9.6 antibody has much higher affinity to RNA:DNA hybrids than to the double-strand RNAs, and the minimum length of the hybrids should be more than 8 base-pair. The structure of Fab binding to the hybrids suggested one loop of S9.6 heavy chain inserts into the minor groove in the RNA:DNA hybrids, and the other three loops flank to the hybrids. The top of four loop all enrich with aromatic or positive charge residues which are potentially responsible for nucleic acids binding specificity. Our results revealed the recognition mechanism of S9.6 on R-loops, which directs the future engineering of S9.6, and thus could further promote R-loop biology studies.

---

R-loops are chromatin structures consisting of one RNA:DNA hybrid strand and the other single stranded DNA (ssDNA) strand (Santos-Pereira and Aguilera, 2015). R-loops are considered to be usually co-transcriptionally formed in cis with the newly synthesized RNA behind the RNA polymerase, and many essential biological functions had been uncovered (Costantino and Koshland, 2015; Crossley et al., 2019; Garcia-Muse and Aguilera, 2019; Niehrs and Luke, 2020; Zhou et al., 2022). In the last decade, a number of genome-wide R-loop mapping techniques had been developed (Crossley et al., 2019; Vanoosthuyse, 2018; Zhou et al., 2022). Among these R-loop profiling techniques, the monoclonal antibody S9.6 had been widely used to detect the RNA:DNA hybrid stand in the R-loops (Chen et al., 2017; Crossley et al., 2020; Dumelie and Jaffrey, 2017; Dutrow et al., 2008; El Hage et al., 2014; Ginno et al., 2012; Hu et al., 2006; Li et al., 2020; Sanz et al., 2016; Wahba et al., 2016; Xu et al., 2020; Xu et al., 2017; Yan et al., 2020). However, it had been shown that the S9.6 could also bind double-strand RNAs (dsRNAs), even with much lower affinity comparing to RNA:DNA hybrids ((Phillips et al., 2013), this study), which challenges the application of S9.6 in R-loop detection (Hartono et al., 2018; Konig et al., 2017). Meanwhile, it is unclear whether the binding of S9.6 to RNA:DNA hybrids has inherent sequence specificity, or the S9.6 prefers the specific structures of substrates, as different double-strand nucleic acids have distinctive features (Liu et al., 2019). Thus, it is worth to reveal the recognition basis of S9.6 to the nucleic acid substrates. The cryo-electron microscopy (cryo-EM) single particle analysis (SPA) technology can directly visualize three-dimensional (3D) structures of bio-macromolecules close to native state at increasingly higher resolution without crystallization (Cheng, 2018; Cheng et al., 2015). In this study, we have applied cryo-EM to analyze the structure of antigen-binding fragment (Fab) of S9.6 bound with RNA:DNA hybrids, the findings give us insight into the recognition mechanism of S9.6 and provide theoretical basis for the application and optimization of R-loop detections.

Initially, we used two ways to obtain the monoclonal S9.6 antibody from the hybridoma cell HB-8730 (ATCC), one is from harvesting cell supernatant by cell culture, the other is the ascites production (Fig S1A). After affinity chromatography, the pure S9.6 antibody were obtained (Fig S1B). Slot Blot results showed that S9.6 derived from both supernatant and ascites fluids have potent R-loop recognition ability (Fig S1C). The S9.6 Fab fragments were generated efficiently from S9.6 IgG after papain digestion (Fig S1A). After affinity chromatography, ion exchange chromatography and size exclusion chromatography, pure Fab was obtained as only the peak and adjacent samples were retained at each chromatography (Fig S1D-H). As cryo-EM requires much less protein than crystallization (Cheng, 2018), we then used the purified Fab for negative staining, and the results showed the S9.6 Fab was uniformly distributed and free of impurities, indicating the Fab is practicable for cyro-EM observation. We synthesized 40 base-pair (bp) long RNA:DNA substrates to observe nucleic acid filaments under cryo-EM, the sequence of the nucleic acid substrates was randomly selected from the ssDRIP (single-strand DNA:RNA immunoprecipitation by S9.6, followed by sequencing) sequencing data (Xu et al., 2017), and the GC content (%) is 50 (Table S1). The DNA or RNA oligonucleotides were labelled with FAM dyes at the 5′-end.

To evaluate the quality and binding ability of the S9.6 Fab and compare the affinity with intact S9.6 antibody, we applied electromobility shift assays (EMSA) and microscale thermophoresis (MST). Indeed, we detected electromobility shift products of S9.6 Fab bound with 40-bp RNA:DNA hybrids (RD in short, Fig 1A). The MST data showed a binding curve and provided the K_d_ of 183 (± 4.5) nM (Fig1B-C). Meanwhile, we assessed the affinity of the same quantity of S9.6 at the same conditions, and the results indicated that there are more shifts at the same quantity, and S9.6 showed higher affinity than Fab with a K_d_ of 72.6 (± 8.7) nM (Fig 1D-F). Both the K_d_ of S9.6 Fab and S9.6 are lower to the very recently results, in which the K_d_ of the purified Fab is 232 nM (Bou-Nader et al., 2022). This could most possibly due to the variations in protein sources, methods, hybrids sequence and length, and buffer conditions. Especially, the Fab protein in (Bou-Nader et al., 2022) was expressed in CHO cells and purified, different to the way we generated Fab, which maximumly maintains the native states of the Fab in the S9.6 antibody (Fig S1A).

We then tested the binding ability of S9.6 Fab to 40-bp dsRNA (RR, RNA:RNA duplex) containing same sequence as 40-bp RD by EMSA (Table S1). The quantity of protein and RR substrate is the same to that used in affinity test for DR mentioned above (Fig 1A and 1D), whereas S9.6 Fab and S9.6 bind much less RR than DR at the same condition, especially when using S9.6 Fab (Fig1G-H). Accordingly, we proceeded the competition experiments between the FAM-labeled RD and Cy3-labeled RR, the S9.6 Fab or S9.6 was mixed with two substrates together and incubated at room temperature for 10 minutes before EMSA detection, gradually increasing the amount of RR to 8 times of RD had no effect to RD binding of Fab or S9.6 (Fig1I-J). We then used RNA:DNA hybrids in the length range from 8 to 14bp with 50% GC-content to determine the minimal R-loop binding length of S9.6 Fab, the EMSA results exhibited no binding shifts when RD length was reduced to 8-bp, it is worth mentioning that the longer RD, the more shifts signals (Fig1K), suggesting that longer nucleic acids are conducive to stable binding of S9.6 Fab and S9.6. Moreover, we found that the higher the concentration of salt ions, the weaker the Fab binding ability, but it did not affect the affinity of S9.6 (data not shown).

**Fig 1.**
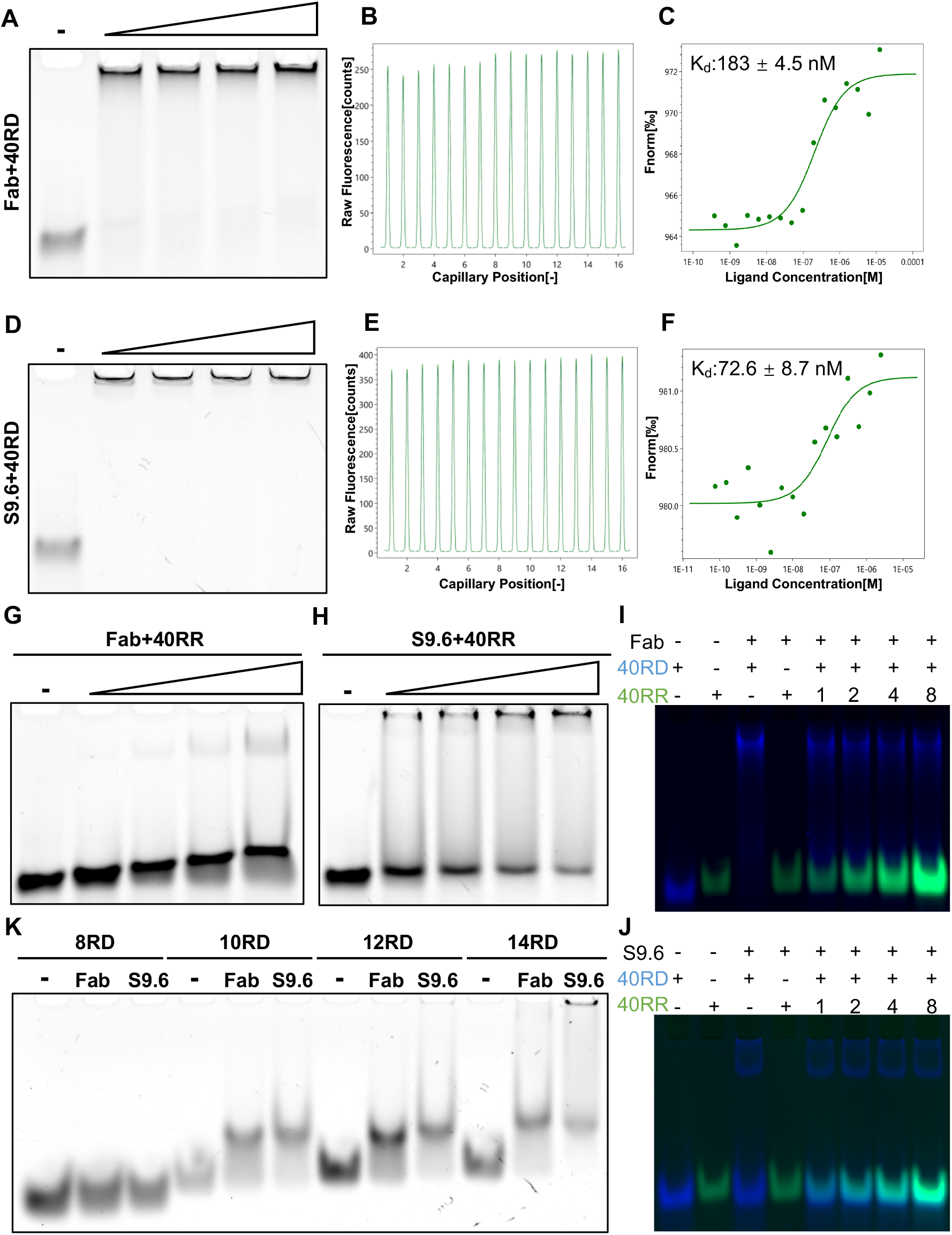
Affinity analysis of S9.6 Fab and S9.6 to RNA:DNA hybrid and dsRNA. (A-C) S9.6 Fab recognizes and binds RNA:DNA hybrid. (A) EMSA detected the binding ability of S9.6 Fab to 40-bp length RNA:DNA hybrid (RD). The DNA was labelled by FAM dyes, 100pM RD was incubated with increasing concentrations of Fab from 2pM to 8pM. (B-C) MST analysis of S9.6 Fab interacts with 40-bp RD. (B) There are no potential sticking effects and fluorescence signals changing during MST progress. (C) The normalized fluorescence of the MST traces was plotted and fitted to calculate the K_d_, the corresponding affinity of Fab with 40-bp RD is 183 nM (± 4.5). (D-F) S9.6 recognizes and binds RNA:DNA hybrid. (D) EMSA detected the binding ability of S9.6 antibody to the same 40-bp length RD, 100pM RD was incubated with increasing concentrations of S9.6 from 2pM to 8pM. (E-F) MST analysis of S9.6 antibody interacts with the 40-bp RD. (E) There are no potential sticking effects and fluorescence signals changing during MST progress. (F)The corresponding K_d_ of S9.6 antibody with the 40-bp RD is 72.6 nM (± 8.7) which indicates higher affinity than S9.6 Fab. (G-H) Both S9.6 Fab and S9.6 bind dsRNA. EMSA detected the binding ability of S9.6 Fab (G) and S9.6 antibody (H) to 40-bp length dsRNA (RR) having the same sequence as 40-bp RD, the RNA was labelled by Cy3 dyes,100pM RR was incubated with increasing concentrations of Fab (G) or S9.6 (H) from 2pM to 8pM. The affinity of the both protein especially Fab to RR is weaker than RD. (I-J) S9.6 Fab and S9.6 preferentially bind RNA:DNA hybrid than dsRNA. Competition experiments of the 100pM 40-bp RD (colored as blue) bound to 2pM S9.6 Fab (I) or 1 pM S9.6 (J) with increasing concentrations of the 40-bp RR (colored as green) from 100 to 800 pM, the RR can’t compete with the RD at this concentration. (K) The affinity of S9.6 Fab and S9.6 to various epitope lengths. EMSA detected the binding ability of S9.6 Fab and S9.6 antibody to different length RD, 2pM protein was incubated with increasing length RD from 8-bp to 14bp, no upper shifts form for the 8-bp RD.

Recently, several macromolecular complexes smaller than 100 kDa have been resolved to high resolution using cryo-EM (Fan et al., 2019; Herzik et al., 2019; Merk et al., 2016). A transmission electron microscope operating at 200keV equipped with a K2 Summit direct electron detector (DED) has been reported resolving the structure of small proteins to high resolution (Herzik et al., 2019). In this study, we further used a transmission electron microscope at 300keV equipped with a K2 DED and GIF Bio-Quantum Energy Filters to resolve the structure of S9.6 Fab complexed with RNA:DNA hybrids which is as small as ∼ 50 kDa. After optimizing sample preparation, the sample showed sufficiently thin ice, which could provide a significantly high signal to noise. S9.6 Fab could be seen directly under micrographs and they appeared to be arranged in a string with low resolution (Fig 2A and 2C). To further understand S9.6 Fab bound to the RNA:DNA hybrids, the specimen was examined under volta phase plate (VPP) with phase shift ranging from 30 to 120 degrees. The specimen demonstrated S9.6 Fab arranged in a string binding to DR with high contrast that was clearly identified, indicating the length of epitope is short and several S9.6 Fab may bind to the same 40-bp RNA:DNA hybrid. This result is in concert with the data shown Fig1K, as more Fab binds to the same RD, more shifts signals could be observed. Moreover, we observed that the higher the ratio of S9.6 Fab to nucleic acids, the more density of the Fab bound to hybrids that would impede particle picking (Fig 2D). We found the binding ability of S9.6 Fab increased with the decreasing NaCl concentration when using EMSA to detect the Fab affinity (data not shown). Consequently, we used a lower concentration of NaCl and relatively lower ratio of Fab to RNA:DNA hybrids to achieve suitable binding conditions for cryo-EM data collection (Fig.2B). Accordingly, we confirmed S9.6 Fab combined with RNA:DNA hybrid by cyro-EM and collected micrographs using conventional imaging approaches without VPP.

**Fig 2.**
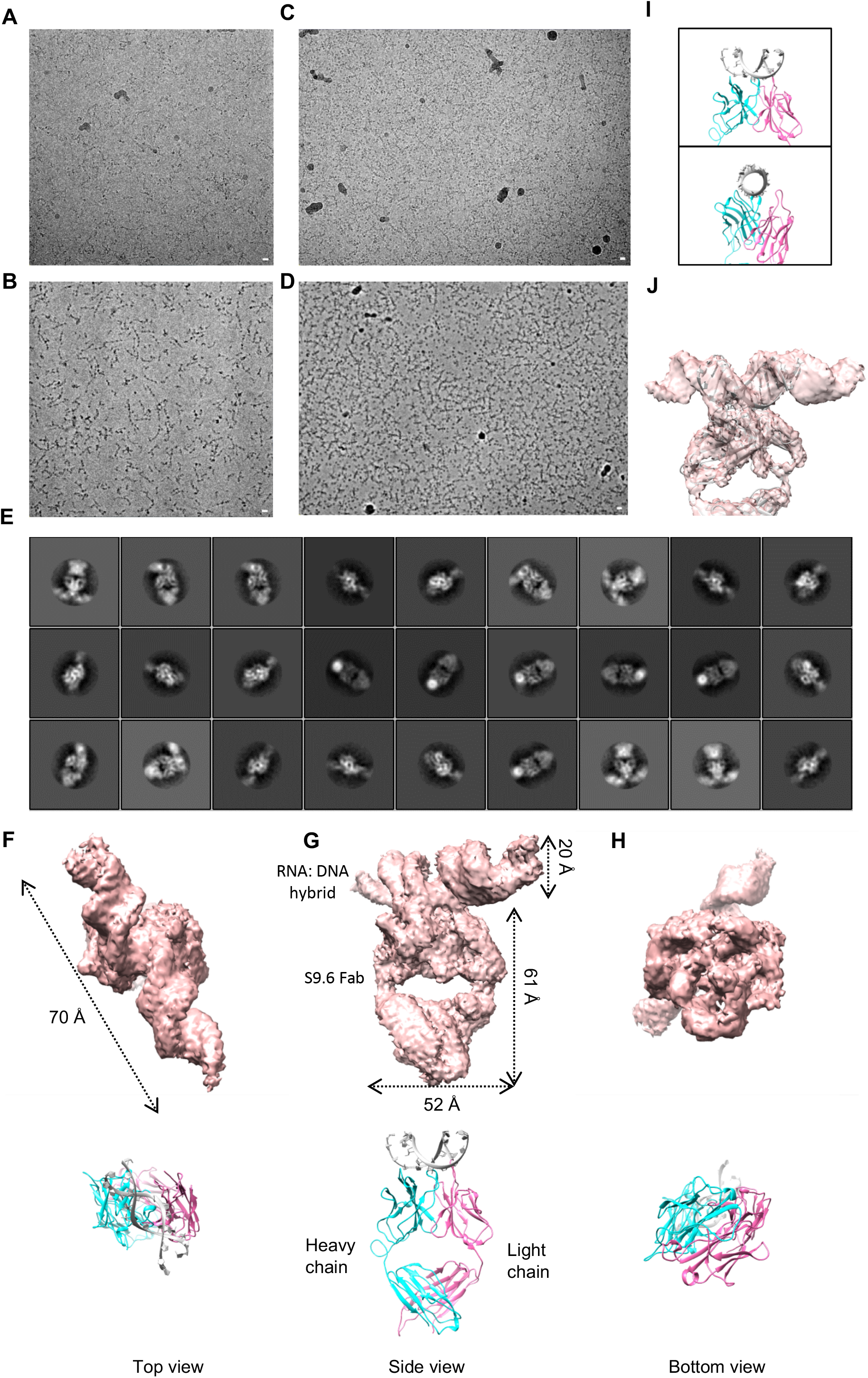
Determination of the cryo-EM structure of S9.6 Fab binding to RNA:DNA hybrid. (A) A representative cryo-electron micrograph of S9.6 Fab to RNA:DNA hybrid solubilized in 0 mM NaCl. (B) A representative cryo-electron micrograph of S9.6 Fab to RNA:DNA hybrid solubilized in 0 mM NaCl by VPP. (C) A representative cryo-electron micrograph of S9.6 Fab to RNA:DNA hybrid solubilized in 50 mM NaCl. (D) A representative cryo-electron micrograph of S9.6 Fab to RNA:DNA hybrid solubilized in 50 mM NaCl by VPP. (E) Representative class averages of S9.6 Fab to RNA:DNA hybrid particles, showing filaments or dot-like RD. (F-H) Top, the indicated view of the sharpened map with a resolution of 4.9 Å. Bottom, predicted structure and cartoon schematic of S9.6 Fab to RNA: DNA hybrid. Heavy chain is in blue, light chain is in pink, RNA: DNA hybrid is in grey. Scale bar, 10 nm. (I)Two views of the S9.6 Fab interface with the RNA: DNA hybrid. (J). The sharpened map fitted with the model 7TQA.

During data processing (Fig S2), 738,447 particles were extracted from high-quality micrographs and applied a 120 Å Fourier high-pass filter prior further processing. The high-pass filter turned out to be necessary for correct alignment of particles. Two-dimensional (2D) alignment and classification from such dataset yielded 2D class averages with clear secondary structural features (Fig 2E). Multiple rounds of three dimensional (3D) classifications using different references were performed to screen the best particles (Fig S2). Finally, we obtained a reconstruction of S9.6 Fab complexed with RNA:DNA hybrids at 4.9 Å resolution with 64,305 particles (Fig S1 J-M and Fig S2). The reconstructed structure of S9.6 Fab bound to RNA:DNA hybrids displayed that the S9.6 Fab is attached to the minor groove of RNA:DNA hybrids, along the spiral direction of the hybrids (Fig2F-H). With this data, we then built and refined a structure model for S9.6 Fab bound to RNA:DNA based on the predicted results of Pyre2 (Kelley et al., 2015). Consistent with the reported crystal structure of IgG Fab (Kaufmann et al., 2002), the S9.6 Fab possesses a hollow ellipsoid-shape architecture that comprises the heavy chain and light chain (Fig 2G). The diameter from the top view and the axial height of the structure are 52 Å and 61 Å, respectively. The diameter and the length of the RNA:DNA are 20 Å and 70 Å, respectively (Fig 2G and 2F). According to our predicted model, two loops of heavy chain and one loop of light chain flank on the RNA:DNA hybrids, and another loop of heavy chain accesses into the inner groove of hybrid (Fig 2I). All of the loops are rich in aromatic amino acids, and the loop extending into the inner groove has an additional positively charged amino acid. Accordingly, we speculated that the loops, deeply inserted into the minor groove may play a key role in the recognition of the RNA:DNA hybrids by aromatic acids and positively charged amino acids. The flanking loops might serve mostly ancillary roles in the recognition process. Collecting more microscopy data should improve the current resolution and could help us confirm our conjecture.

In summary, the native S9.6 antibody derived from hybridoma cells are relatively easy to obtain, from both the cell culture and ascites production (Fig S1A). Both of the S9.6 Fab and S9.6 intact antibody prefer to bind RNA:DNA hybrids than dsRNAs, and the affinity of S9.6 Fab is little weaker than S9.6. Longer length of RNA:DNA hybrids could promote more stable binding of S9.6 and S9.6 Fab, while no binding could be detected by EMSA for the 8-bp RNA:DNA hybrids. We demonstrated that transmission electron microscope at 300keV equipped with a K2 DED and GIF Bio-Quantum Energy Filters could be used to resolve the ∼ 50 kDa protein. S9.6 Fab complexed with RNA:DNA hybrids is resolved to 4.9 Å resolution and is expected to higher resolution if more data are collected. The current resolution structure exhibits that three loops formed by the variable region of Fab heavy chain and one loop formed by the variable region of Fab light chain are close to the minor groove of hybrids. One loop of the heavy chain embeds into the inner groove and carries both aromatic and positively charged amino acids that may play a major role in recognition of S9.6 Fab to RNA:DNA hybrids, and the other three loop that all have aromatic amino acids may help recognition.

R-loops have both positive and negative functions in genome regulation and maintenance (Crossley et al., 2019; Garcia-Muse and Aguilera, 2019; Niehrs and Luke, 2020; Zhou et al., 2022). The S9.6 antibody that specifically binds to RNA:DNA hybrids is the most widely used tool in characterizing R-loops. In this study, we tested the binding ability of native S9.6 Fab and S9.6 by EMSA and MST in vitro, the Fab showed slightly weaker affinity to RNA:DNA hybrids when comparing to the intact S9.6 antibody. We speculated that the Fab region recognizing DNA strands of hybrids becomes more flexible without Fc fragment, that implies the hinge and Fc fragment in the S9.6 antibody might enhance the binding affinity of S9.6 Fab. Besides, the minimum length of the hybrids recognized by native S9.6 should be longer than 8-bp in vitro binding experiment, even though the epitope length determined by structure analysis is 6-bp and the single-chain variable fragment (scFv) of S9.6 can bind to 6-bp hybrids (Phillips et al., 2013). This might be because S9.6 antibody Fab linked by disulfide bonds is more flexible and could not recognize the secondary structure formed by only 8-bp and shorter hybrids, whereas scFv is fused together by repeating protein linker (Phillips et al., 2013). Although the non-specific dsRNA recognition of S9.6 questioned the usage of S9.6 in R-loop mapping, we found that S9.6 prefers RNA:DNA but could not bind dsRNAs when RNA:DNA and dsRNA are both present in the reaction (Fig 1I and 1J).

Notably, cryo-EM enables 3D structure determination of biological specimens in vitrified state without crystallization, which has not only greatly increased the throughput of high-resolution structure determination, but has also allowed for the 3D visualization of macromolecular complexes previously deemed intractable for structural studies due to size, conformational heterogeneity, and compositional variability. Despite these advances, it is currently difficult to resolve the structure of small proteins (<100 Kda) using Cryo-EM mainly due to the low signal-to-noise. While the VPP has enabled visualization of specimens in this size range, this instrumentation is not fully automated and can present technical challenge. Here, we showed that conventional 300kV microscopes equipped with GIF can be used to determine the structure of small protein ∼50 kDa without symmetry. We resolved the structure of ∼50 kDa S9.6 Fab bound with RNA:DNA hybrids at 4.9 Å resolution using only 64,305 particles, and higher resolution could be achieved once enough data were collected.

## Figure legend

**Fig S1.**
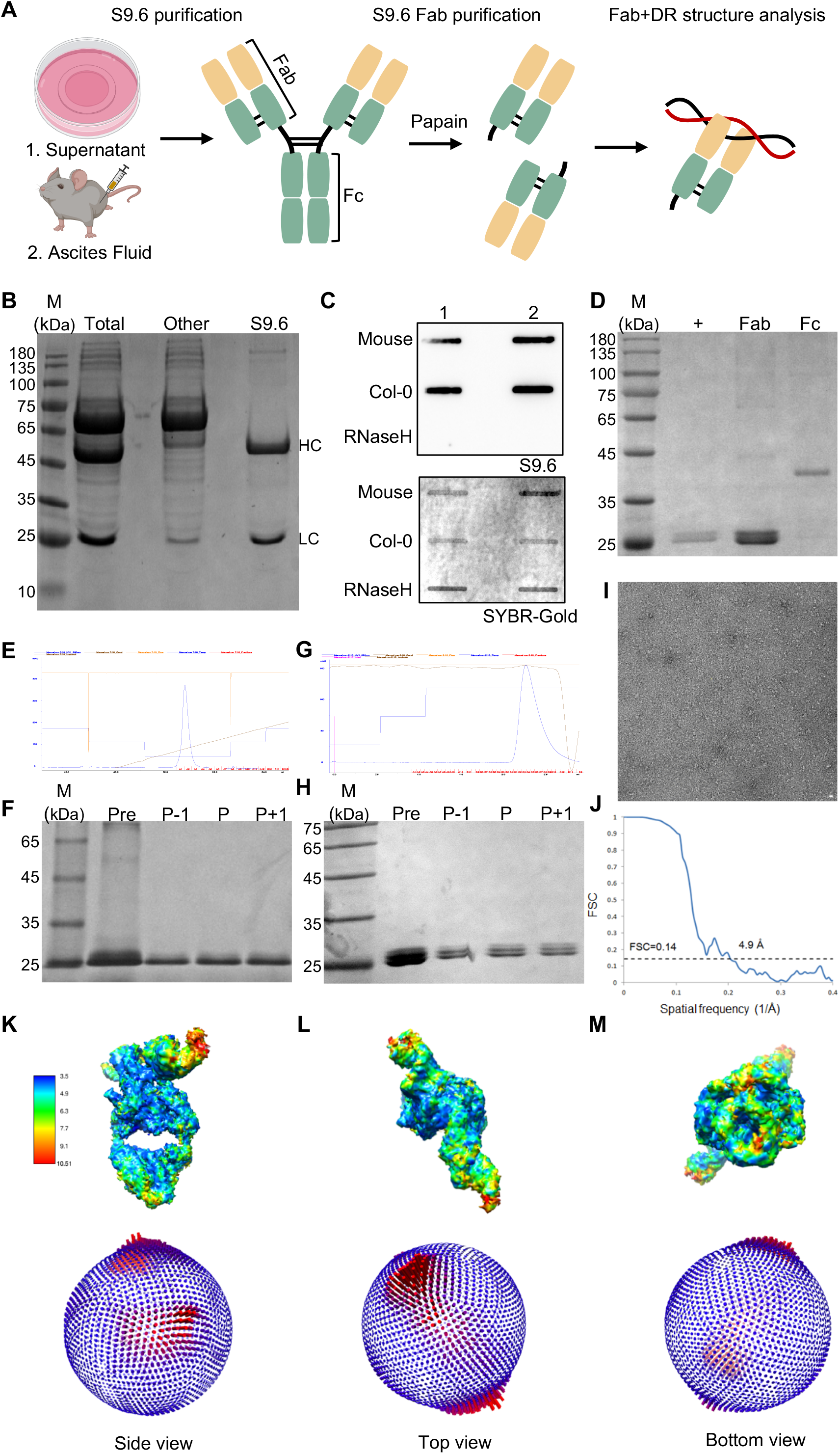
Purification and preparation of S9.6 Fab for Cryo-EM. (A) Schematic diagram of S9.6 and S9.6 Fab prepared for Cyro-EM analysis. Monoclonal S9.6 antibody is produced from hybridoma cell HB8730, the cells were cultured both in medium and peritoneal cavity of mice, the purified S9.6 from cell supernatant or ascites fluid then treated with papain enzymes to obtain Fab regions of S9.6. Fab complexed with 40-bp RD were analyzed for Cyro-EM structure. (B) Affinity chromatography purification of S9.6. Denaturing SDS-PAGE analysis of primitive supernatant (Total), purified S9.6 (S9.6) and miscellaneous protein (Other), most miscellaneous proteins were removed after affinity purification. M, protein marker, HC, S9.6 heavy chain, LC, S9.6 light chain. (C) Slot Blot detection of S9.6 recognizing ability for R-loops. 100 ng genomic DNA of mouse embryonic fibroblast cells and Arabidopsis wild-type seedlings were detected by S9.6 from supernatant (1) and ascites (2) respectively, RNase H is the negative control of Arabidopsis genomic DNA digested with commercial RNase H. SYBR-Gold stain and indicate genomic DNA. (D-H) Purification of S9.6 Fab. (D) S9.6 Fab was first purified with HiTrap Protein A columns after digested by papain, S9.6 Fab and Fc were completely separated. +, Control. (E-F) S9.6 Fab obtained from (D) was further purified through ion exchange chromatography using Source S colum ns. The remaining impurities was washed away at low ion concentrations, S9.6 Fab was eluted at nearly 100 mM NaCl concentration showed in (E). (F) Denaturing SDS-PAGE analysis of S9.6 Fab elution. Pre, pre-purification sample, P-1, Fab from tube in front of elution peak, P, Fab from tube of elution peak, P+1, Fab from tube behind the elution peak. (G-H) S9.6 Fab obtained from (E) was finally purified with size exclusion chromatography. (G) S9.6 Fab was collected near the corresponding position of 50 kDa. (H) Denaturing SDS-PAGE analysis of (G). Pre, pre-purification sample, P-1, Fab from tube in front of elution peak, P, Fab from tube of elution peak, P+1, Fab from tube behind the elution peak. (I) A representative negative-staining electron microscopy image of S9.6 Fab to RNA:DNA hybrid solubilized in 0 mM NaCl. (J) Gold standard Fourier shell correlation (FSC) curves of the indicated density map. Reported resolution was based on the FSC = 0.143 criterium. (K-M) Top, local-resolution maps of the indicated densities. Bottom, Euler angle distribution of particles that were used in the final 3D reconstruction; the height of the cylinder is proportion to the number of particles for that view. Scale bar, 10 nm.

**Fig S2.**
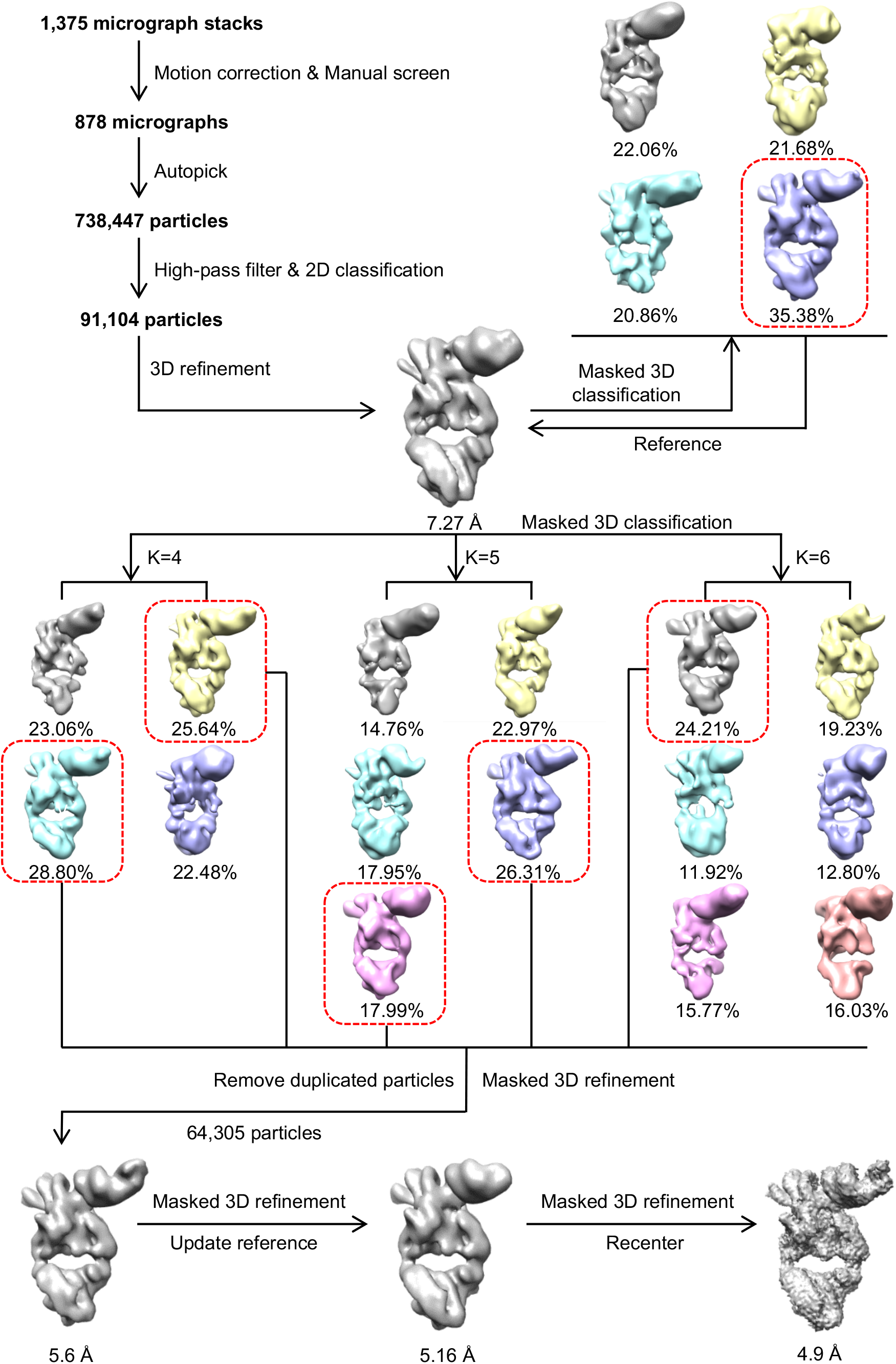
Flowchart of electron-microscopy data processing. Details of data processing are described in the ‘Image processing’ section of the Methods.

## Methods

### Generation and purification of S9.6

S9.6 hybridoma cell, HB-8730™, was purchased from ATCC (American Type Culture Collection), cells were cultured in complete medium (DMEM Dulbecco’s Modified Eagle’s Medium, GIBCO) supplemented with 10% fetal bovine serum (GIBCO) and 1% penicillin/streptomycin. The cell culture was firstly incubated at 37°C with 5% CO2 in incubator, then the cells were transferred into 5L culture flask for large-scale culture and proceeded mice celiac inoculation at the same time. Harvest the cell culture supernatant or ascites fluid that contain S9.6 antibody. The supernatant was firstly concentrated to 200 mL using an ultrafiltration membrane of 50 KDa, then the concentrated supernatant or ascites fluid was purified separately by affinity chromatography with HiTrap Protein G HP columns (5mL). S9.6 from supernatant needs further ion exchange purification because of more impure protein left. Finally, the eluted S9.6 antibody was divided into 4mg per tube for lyophilization after desalting and was stored in -80 □.

### Generation and purification of S9.6 Fab

Dissolved 4mg of the S9.6 powder in 0.5 mL digestion buffer (200mM Hepes,20 mM cysteine-HCl, PH 7.0), added 0.5 mL immobilized papain slurry after equilibrated with digestion buffer, incubated in a shake incubator for 5 hours at 37 □. Separated the immobilized papain from the digest using a resin separator, purified the S9.6 Fab from IgG and Fc fragments using HiTrap Protein A HP columns (5mL). After ion exchange chromatography, the protein is further purified by size exclusion chromatography and desalted.

### Synthesis of nucleic acids

The RNA and DNA oligonucleotides were synthesized by the GenScript Biotech Corporation, the sequences are showed in Table S1. For RNA: DNA hybrids, 5′-end of DNA was labelled with FAM, and for dsRNA, the 5′-end of RNA having the same sequence with DNA was labelled with Cy3. Dissolved and mixed equal molar of oligonucleotides in annealing buffer (100 mM KCl; 10 mM Hepes, pH 7.2) and incubated at 95 □ for 1 minute,75 □ for 15 minute, and then slowly cooled down to room temperature(Hua et al., 2018). The same nucleic acids were used in EMSA and MST.

### Electromobility shift assay (EMSA)

100 pM RNA: DNA hybrids or dsRNAs were mixed with increasing amount of S9.6 Fab or S9.6 in 10 μL reaction buffer (20 mM Hepes, 2.5 mM MgCl_2_, 2.5% Glycerol), the mixture was incubated in dark for 10 minutes at room temperature. 10 μL products were loaded into 7.5% native-PAGE gels in 0.5 x TBE and separated at 10V/cm, the gel was visualized using the Typhoon FLA 9500 Instrument (GE Healthcare Life Sciences).

### Microscale thermophoresis (MST)

The Monolith NT.115 from NanoTemper Technologies and Monolith NT capillaries (Standard treated, NanoTemper Technologies) were used for MST, the procedures mainly followed the manuscript of the User Starting Guide Monolith NT.115 (Jerabek-Willemsen et al., 2014). In our MST reaction system, reaction and dilution buffer were MST buffer with little modified (50mM Tris-HCl pH7.6, 50mM NaCl, 10mM MgCl2, 0,05% Tween-20), 50 nM S9.6 Fab or S9.6 were used as starting concentration and 1:1 diluted, and 12.5 μM RNA: DNA hybrids were used totally, 40 % MST power was used after auto-detection. The MO.Affinity Analysis Software was used to analyze the recorded MST signal and derive a binding curve from the thermophoresis traces.Details

### Negative-staining electron microscopy

4 ul freshly prepared sample was applied to a glow-discharged (PLECO) carbon coated copper grid (300 mesh, Zhongjingkeyi Technology). After the grid was incubated at room temperature for 1 min, the excess liquid was removed with filter paper, and then the grid was stained by a drop of 2% uranyl acetate for 20 s. The staining process was repeated twice and the third staining process was carried out additionally 1 min. Finally, the excess staining buffer was removed and air-dried. The prepared grids were observed on a T12 microscope (FEI) operating at 120 kV, using a 4k x 4k CCD camera (UltraScan, Gatan) at a nominal magnification of 49,000 with a calibrated pixel size of 2.3 Å.

### Electron microscopy sample preparation and image collection

The cryo-samples were prepared using a Vitrobot Mark IV (Thermo Fisher) operated at 8 °C and 100% humidity. In brief, 4 ul freshly prepared sample was applied to a glow-discharged 300 mesh Quantifoil Au R2/1 grid. The grid was blotted for 1 s with a nominal blot force 0 of the Vitrobot without incubation. And then the grid was immediately plunged into pre-cooled liquid ethane for vitrification.

The prepared grids were observed using a 300 kV Titan Krios (Thermo Fisher) electron microscope equipped with a Cs corrector and GIF Quantum energy filter (slit width 20 eV). Images were recorded by a K2 Summit direct electron detector working at the super resolution mode. Data collection was performed using AutoEMation [1] with a nominal magnification of 105,000 yielding a super-resolution pixel size of 0.5455 Å, and with a defocus range from -1.5 to -2.4 μm. Each micrograph stack, which contained 32 frames, was exposed for 5.6 s with a total dose of 50 e^−^ /Å^2^. The VPP data collection was performed according to following protocol

### [5].Cryo-EM image processing

The procedure for image processing of S9.6 Fab to RNA: DNA hybrid is presented in fig.S2. A total of 1,375 micrographs were collected. All the micrographs were motion-corrected using MotionCor2 [2] with a bining factor of 2, resulting in a pixel size of 1.091 Å^2^. Contast transfer function (CTF) parameters were estimated by CTFFIND4.1 [3]. A total of 738,447 particles were automatically picked by Relion [4]. After particles were extracted by Relion, a120 Å high pass filter was applied to the particle stacks for a better 2D classification performance. Finally, 91,104 particles were selected for the further 3D refinement. The 3D structure of negative stain was used as initial model for the first round of 3D refinement. After that, the map was used as reference for 3D classification. The best class of 3D classification was used as updated reference for three parallel masked 3D classifications, which were performed with three, four and five classes, respectively. Good particles from each class were merged, removing duplicated particles, and subjected to 3D refinement, resulting in a map at 5.6 Å. Then each particle was recentered and re-extracted without bining from motion-corrected integrated images. After 3D refinement, the dataset yielded a map at 4.9 Å resolution. The reported resolution was based on the gold-standard Fourier shell correlation at 0.143 criteria. Variations in local resolution were estimated usign Resmap [6].

## Acknowledgements

We thank Dr. Jun-Jie G. Liu and S. Fan for the useful discussions and suggestions; H. Wang, N. Liu and J. Xu for providing the graphene grids; C. Yan for sharing the scripts for deep-2D; J. Lei and X. Li for technical help. We thank the Protein Preparation and Identification Facility at Technology Center for Protein Science in Tsinghua University for purification of S9.6; the Center of Biomedical Analysis in Tsinghua University for affinity detection of S9.6; the Beijing Advanced Innovation Center for Structural Biology for facility and financial support.

## Author contributions

Q. Sun, X. Li and H. Li conceived the study and designed the experiments with Q. Li and C. Lin; <u>Q. Li performed</u> protein purification, affinity detection, sample preparation for cryo-EM and partial cryo-EM data collection; C. Lin performed cryo-EM data collection and analysis.

